# Engineered tRNA suppression of a CFTR nonsense mutation

**DOI:** 10.1101/088690

**Authors:** John D. Lueck, Daniel T. Infield, Adam L. Mackey, R. Marshall Pope, Paul B. McCray, Christopher A. Ahern

**Affiliations:** Department of Molecular Physiology and Biophysics, University of Iowa, Iowa City, IA, 52242, USA.

## Abstract

Ten percent of human diseases are caused by ‘nonsense’ mutations that lead to premature truncation of the protein reading frame. Small molecules that promote read-through of such PTC have significant clinical promise but current iterations suffer from low *in vivo* efficacy and the nonselective amino acid incorporation. Alternatively, while gene-modifying approaches, such as CRISPR/Cas9, represent a long-term solution, such treatments may be far from reaching the clinical setting. Building on previous work by our group and others, we describe a tRNA engineering approach that enables the conversion of an in frame nonsense stop mutation to the naturally occurring amino acid, thus rescuing the full-length wild type protein. Data is presented demonstrating the functionality of the approach with the rescue of CFTR W1282X, a human mutation that causes cystic fibrosis (CF). The stringency of the approach is confirmed by mass spectrometry in a model protein indicating the encoding of only tryptophan at the TGA suppression site. The data describe the first use of an edited tRNA to repair a CF causative mutation and serve a proof of principle for the eventual use of codon-edited tRNA for the therapeutic rescue of PTC disease codons.

## INTRODUCTION

Nonsense disease causing mutations result from nucleotide changes in the reading frame of a protein that encode stop codons (TAG, TGA or TAA). Such genetic alterations in the reading frame of cystic fibrosis transmembrane conductance regulator (CFTR) are responsible for upwards of ten percent of cystic fibrosis cases (Rowe, Miller et al. 2005). An example of one such mutation is W1282X, a premature termination codon (PTC) which causes a loss of CFTR function and produces severe cystic fibrosis (CF) phenotypes (Rowe, Miller et al. 2005). Of relevance for the therapeutic management of CFTR PTC mutations, small molecules have been identified which promote read-through of disease producing nonsense mutations (Bedwell, Kaenjak et al. 1997, Keeling, Xue et al. 2014, Mutyam, Du et al. 2016). However, these approaches have a number of challenges yet to be overcome. The type of amino acid that replaces the stop codon is difficult to control in mammalian cells, often leading to a missense mutation at the site of the original PTC (Roy, Friesen et al. 2016). Therefore, when therapeutically assisted PTC stop codon read-through is successful, the non-selective incorporation of an amino acid at the location of the nonsense codon has the potential to affect protein folding, trafficking and function; and thus requires additional therapeutic intervention (Xue, Mutyam et al. 2014). Further, some such compounds have unexpectedly low efficiency of codon skipping *in vivo (Zomer-van Ommen, Vijftigschild et al. 2016).*These issues are likely to be compounded by the general reduction in mRNA abundance of CFTR PTC message (Hamosh, Rosenstein et al. 1992). The widespread use of these compounds to repair PTC *in vivo*(Kerem, Konstan et al. 2014) and the unexpected discovery that endogenous stop codon read-through is common in animals (Jungreis, Lin et al. 2011), suggests that assisted suppression at the site of disease PTC could be a viable therapeutic approach if delivered to a subset of cell types, i.e., airway epithelia. Importantly, the available data suggest that there are active biological systems in place which monitor and minimize the impact of stop codon read-through at so-called ‘real’ stop codons that terminate protein synthesis (Bengtson and Joazeiro 2010). Lastly, as an alternative to the use of PTC skipping compounds, the repurposing of the RNA editing complex ADAR for possible permanent repair of the PTC is also an active area of investigation (Montiel-Gonzalez, Vallecillo-Viejo et al. 2013). While methods are being developed for the therapeutic repair of CF PTC disease mutations, there remains an unmet need to for therapeutic suppressors for use in CF and generally, more effective treatments of PTC diseases. Further, redundant experimental efforts on multiple fronts may be a successful strategy for the rescue of CFTR nonsense codons and other such diseases, given that each mutation may yield novel phenotypes based on the position within the gene or other complicating factors.

A general approach is proposed herein where tRNA sequences are edited to no longer recognize and suppress their cognate DNA codon sequences, but instead guide the tRNA (and cognate amino acid) to the disease missense codon PTC, thus fully ‘repairing’ the PTC, resultingin wild type protein sequence. Once identified, the subsequent expression of this tRNA within a target cell will serve to ‘suppress’ the disease PTC upon being acylated (aka ‘charged’) by its cognate synthetase that is endogenously expressed in the cell. The key to the overall success of this strategy relies on the ability of the tRNA to tolerate the mutation of the anticodon, a determinant of synthetase recognition. It has been noted that PTCs not only are a major cause of human disease, but that suppressor tRNA could also be effective for both their identification and as possible therapeutic agents (Capone, Sedivy et al. 1986, Atkinson and Martin 1994). One proof of principle example of this approach has been proposed for the site-specific editing of a tRNA-Lys (a tRNA that encodes for the amino acid lysine) that was repurposed for a TAG (amber) PTC in the mRNA encoding beta-thalassemia (Temple, Dozy et al. 1982). In this example, the engineered lysine tRNA was edited to recognize the PTC amber stop codon and encode lysine, thereby repairing/completing the beta-thalassemia reading frame. It is worth noting that each amino acid, while having a single synthetase, has many tRNA, upwards of 20+ in some examples (Chan 2009) and more than 450 total tRNAs annotated expressed in humans (Lowe and Eddy 1997, Lowe and Chan 2016). Thus, there is a biologically rich source of tRNAs to mine for new tRNA functionality. Similar approaches in fruit fly indicate that the overexpression of suppressor tRNA in multicellular organisms is viable (Laski, Ganguly et al. 1989). This possibility is consistent with the viability of transgenic *Drosophila* and *C. elegans* organisms that over-express amber suppressor tRNA (Bianco, Townsley et al. 2012, Chin 2014). Further, it has been shown that cell lines with stable expression of suppressor orthogonal tRNAs are viable (Koukuntla, Ramsey et al. 2013). There are numerous examples of the use of codon-edited or suppressor tRNAs that when over-expressed in target cells or host-organisms result in biological activity that is not paired with toxicity, suggesting low suppression efficacy at bona fide termination codons, or that such events have limited cellular impact.

We deemed CFTR an attractive target for rescue of PTCs by engineered tRNA, not only because a significant proportion of people with CF harbor PTCs, but also because several lines of evidence suggest that a fraction of restored or residual normal anion channel function is sufficient to eliminate the most devastating symptoms of cystic fibrosis (Sosnay, Siklosi et al. 2013, Char, Wolfe et al. 2014).

## RESULTS

Given the available published data and our previous work with edited tRNAs, we reasoned if it might be possible to express eukaryotic tRNA that had been anticodon edited to suppress stop sites, TGA for instance, in a CFTR cDNA construct. For simplicity, we have termed these engineered tRNA molecules as ACE-tRNA, for their anticodon editing. As a preliminary experimental foray, four of the nine human Trp tRNA were synthesized as TGA suppressors by making the relevant mutations within the anti-codon region of the tRNA, Figure 1. These tRNA were then co-expressed with CFTR Trp1282X channels by transfection in HEK293T cells, Figure 1B. Figure 1C shows that ACE-tRNA_Trp_ #1 and #3 show very modest rescue of the full length CFTR while #2 has no apparent functional effect, possibly due to intolerance of this specific tryptophan tRNA to anti-codon editing. However, ACE-tRNA_Trp_ #4 showed robust rescue of full-length mature CFTR W1282TGA channel. Note that the Trp tRNA is very likely to be acylated by the endogenous tryptophan synthetase, thus one would expect the rescued CFTR channels to have a Trp at W1282. Consistent with this possibility, the expression conditions produced both CFTR B and C bands indicated by arrows, Figure 1C, with the higher band corresponding to the glycosylated mature functional CFTR form.

**Figure 1.**
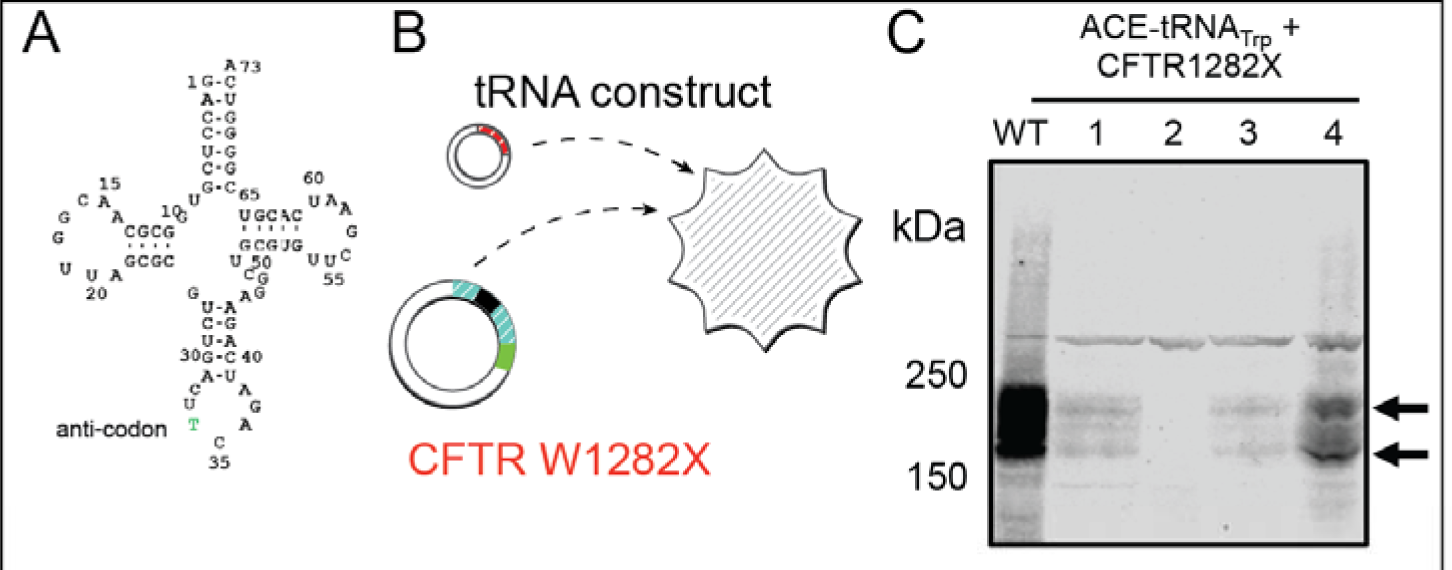
Rescue of CFTR W1282X with ACE-tRNA. A, the human Trptophan. tRNA with anti-codon labled. B, schematic of experimental approach for transient expression of the indicated constructs. C, left panel, 4 of 9 human Trp tRNA were screened for anti-codon editing tolerance and PTC suppression efficiency. ACE-Trp tRNA #4 displayed the best ability to rescue the W1282X PTC in HEK cel ls as demonstrated by Western blot with CFF antibody M3A7. Note the presence of the two bands in the #4 tRNA rescue indicated by arrows indicating the lower 'B' and upper, trafficked and glycosylated 'C' band.

To examine the stringency of the suppression process in higher resolution, and particularly, identification of the amino acid that is being encoded at the TGA site, we used mass spectrometry of a model protein histidinol dehydrogenase (HDH), Figure 2. Here, a TGA codon was introduced at N94 by standard site directed mutagenesis. This construct was then co-expressed with ACE-tRNA_Trp_ #4 from Figure 1, purified and analyzed by mass spectrometry. Search parameters were set initially so as to identify fully tryptic peptides with a precursor mass accuracy of 10 ppm and mass accuracies for MS/MS fragments set to 0.4 for CID/ IT detection and 0.02 Da for HCD activation, wherein fragment ions were detected in the Orbitrap at a mass resolution of 30,000. Searches were tuned to maintain a 1% or lower FDR for protein alignments. These initial searches assumed a possible two missed tryptic cleavages and fixed carbamidomethyl modification for Cys residues (+57.02146), as well as variable (di-) carbamidomethylation for the N terminus of any peptide, possible propionamide adduction at Cys residues (+71.03711) and variable oxidation of Met (+15.99491) residues. These searches were also set to identify the custom modification Asn to Trp (+72.03638).

**Figure 2.**
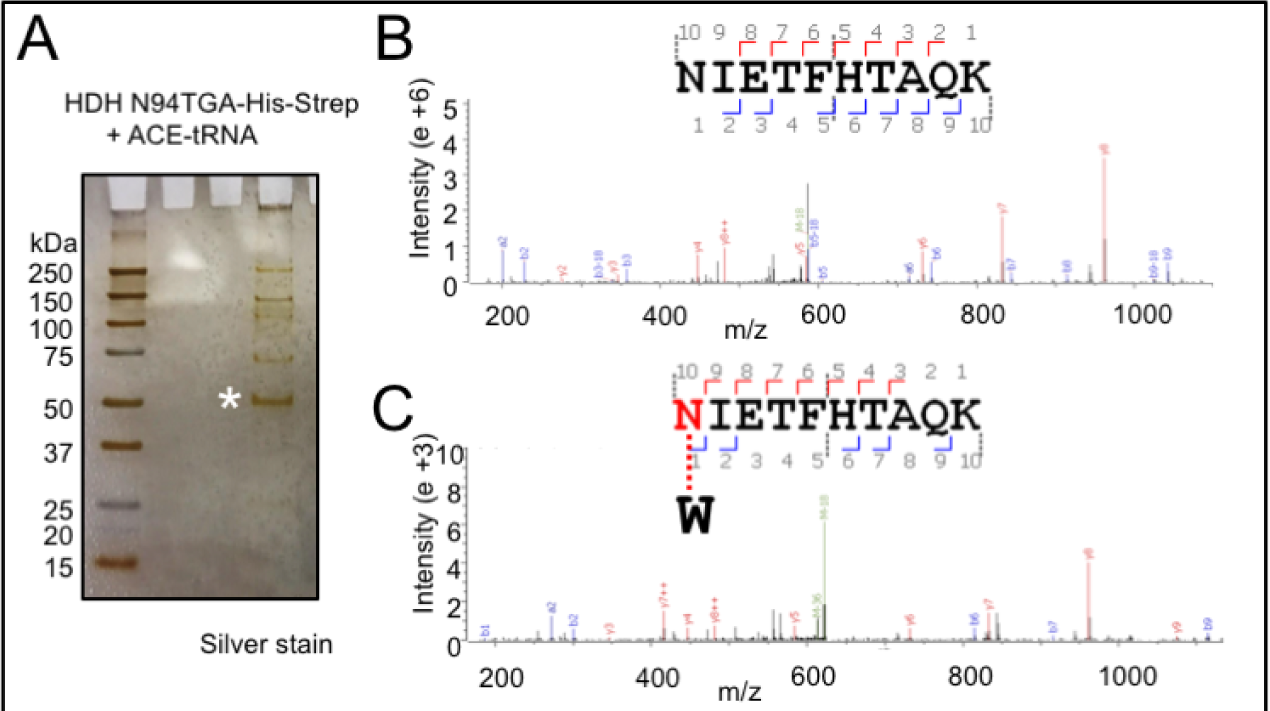
Expression of ACE-tRNAr_Trp_ results in specific incorporation of tryptophan into nonsense stop sites. A) Co-expression of HOH-His-Strep N94TGA and ACE-tRNA #4 results in robust expression of full length HDH protein that is detectable by silver stain. The asterisk indicates full-length HDH protein that was purified from Hek293 cells by StrepTrap affinity chromatography. B) Spectra of WT HDH displays an asparagine at position 94. C) Spectra from HDH N94TGA protein rescued with ACE-tRNA shown in panel A) indicates that position 94 is suppressed with tryptophan. Spectra are representative of >6 peptide spectra.

More detailed searches were designed to identify other alterations against a smaller, focused data base containing the most credentialed 279 sequence alignments and decoys. Specifically, all potential amino acid substitutions that may take place at Asn residues were searched in groups of five. Additional searches screened for “Wild Card” mass shifts between −290 and +290 Th confined to Asn residues. Importantly, the searches identified no alternative substitutions at the HDH N94X site other than the ACE-tRNA_Trp_ encoded tryptophan.

## DISCUSSION

Nonsense mutations are implicated in several human diseases including CF, wherein they are found in approximately 10% of patients. Pharmacological strategies that promote skipping of the stop codon have not thus far been demonstrated to be efficacious in patients (Kerem, Konstan et al. 2014). In the current study, a foundation has been laid to repair nonsense mutations in CFTR and other proteins using an engineered tRNA. To our knowledge, this is the first description of the use of such an approach for the rescue of a CFTR nonsense codon. Rescue has been determined biochemically and functional studies are underway in multiple cell types. Notably, biochemical examination of rescued CFTR W1282X channels indicated the presence of a significant population of glycosylated mature CFTR protein, indicative of plasma membrane expression. We then show by mass spectrometry that at the site of repair of the TGA codon in a model protein is successfully and stringently replaced with a tryptophan side-chain.

The tRNA codon-edited approach offers a number of significant benefits over existing strategies: 1) Improved codon specificity – the expressed ACE-tRNA is directed towards a specific stop codon (TGA, for example, over TAA and TAG), thus reducing off-target effects at stop codons unrelated to disease. 2) Amino-acid specificity – the expressed ACE-tRNA is engineered to transfer the wild-type amino acid encoded by normal CFTR cDNA sequence, thus negating any spurious effects on CFTR stability, folding, and trafficking. 3) Tunability – the system can be personalized for each type of tRNA and PTC mutation. 4) Compact design – the essential tRNA expression cassette is <200 base pairs and can be multiplexed to accommodate as many tRNA repeat quantities needed for tunable expression levels. 5) Proof of principle for a general strategy – in-frame stop codons are a major cause of human disease and few treatment options exist; thus the data suggest that ACE-tRNA may lead to insights for the development of new therapeutics for genome wide disease causing nonsense mutations.

## MATERIALS AND METHODS

### ACE-tRNA, HDH-His-Strep and CFTR Expression Plasmids

The cDNA for the coding region and 200bp of the 3’ UTR of human CFTR was ligated into pcDNA3.1(+) (Promega, USA) using the KpnI and XbaI restriction enzymes. The W1282TGA mutation was introduced using QuickChange XL II (Stratagene, USA). The cDNA encoding the E. coli histidinol dehydrogenase was codon optimized for *mus musculus* and synthesized (Bio Basic Inc, Canada) with a c-terminal 8xHis-Strep-tag for protein purification from mammalian cells. The synthesized cDNA was ligated into pcDNA3.1(+) using EcoRI and XhoI restriction sites. Four tryptophan tRNA genes were chosen from GtRNAdb (Chan and Lowe, 2009). The 5' leader sequence taken from the human tRNA_Tyr_ gene (Ye et al., 2008) was ligated such that a golden gate MCS would position a correct transcription startA 3’ termination sequence (*GTCCTTTTTTTG*) was inserted 3’ of the golden gate MCS such that it would follow last indicated tRNA nucleotide. The tRNA oligos were synthesized (IDT, USA) with the cognate anticodon edited to recognize TGA stop sites and ligated into puc57 between the 5’ leader and termination sequence using standard golden gate cloning techniques.

### Cell Culture, Protein Expression and Western Blot

Hek293T cells (ATCC, USA) were grown in standard grown media containing (% in v/v) 10% FBS (HiClone, USA), 1% Pen Strep, 1 % L-Glut in high glucose DMEM (Gibco, USA) at 37º C, 5% CO2. cDNA was transfected at 75% confluency using standard calcium phosphate methods. Following 36hrs the cells were scraped and pelleted at 3,000g in PBS supplemented with 0.5 ¼g/ml pepstatin, 2.5 ¼g/ml aprotinin, 2.5 ¼g/ml leupeptin, 0.1 mM PMSF, 0.75 mM benzamidine. For CFTR expressing cells, the cell pellet was solubilized in a buffer containing 1% triton, 250mM NaCl, 50mM tris-HCl pH 7.4, and 0.5 ¼g/ml pepstatin, 2.5 ¼g/ml aprotinin, 2.5 ¼g/ml leupeptin, 0.1 mM PMSF, 0.75 mM benzamidine. Equal cell-lysate was loaded on a 3-15% gradient SDS-page in the presence of 1% 2-mercaptoethanol, separated at 55 V O/N and transferred to 0.45 ¼M LF PVDF (Bio-Rad, USA). PVDF was immunoblotted using anti-CFTR antibody M3A7(1:1000; Millipore, USA) in 2% NF milk and imaged on LI-COR Odyssey Imaging System (LI-COR, NE, USA). For HDH-His-Strep expressing cells, the cell pellet was vigorously dounce homogenized in 100mM sucrose, 1mM DTT, 1mM EDTA, 20mM tris-HCl pH 8.0, 0.5 ¼g/ml pepstatin, 2.5 ¼g/ml aprotinin, 2.5 ¼g/ml leupeptin, 0.1 mM PMSF and 0.75 mM benzamidine. The lysate was centrifuged at 100,000g for 30min at 4 º C. The supernatant (soluble cellular protein) was separated on SDS-page as before and was immunoblotted using anti-Strep antibody (1:5000; iba, Germany) in 2% NF milk and imaged on LI-COR Odyssey Imaging System (LI-COR,USA).

### Mass Spectrometry

Fragmentation data on purified HDH-His-Strep protein were obtained at the University of Iowa Proteomics Facility. Briefly, HDH-His-Strep protein from the soluble fraction of the high speed spin was passed through StrepTrap HP columns (GE Healthcare, Sweden) and washed with 5 column volumes of 100mM sucrose, 1mM DTT, 1mM EDTA, 20mM tris-HCl pH 8.0, 0.5 ¼g/ml pepstatin, 2.5 ¼g/ml aprotinin, 2.5 ¼g/ml leupeptin, 0.1 mM PMSF and 0.75 mM benzamidine. The protein was eluted in wash buffer supplemented with 10mM d-desthbiotin and concentrated in 30kDA cutoff Amicon-Ultra filtration columns (Millipore, USA). The concentrated protein was loaded on NuPage 4-12% Bis-Tris precast gels (Invitrogen, USA) and separated at 150V for 1.5 hrs. The gel was stained using a Pierce mass spec compatible silver stain kit (Thermo Scientific, USA).

#### In-gel Trypsin Digestion

For this analysis, 20 μg aliquots each fractions were loaded onto separate gel lanes. A 10 μl solution of mass ladder markers (Sharp pre-stained migration standards, Invitrogen) was loaded onto a separate gel lane to serve as a guide to molecular weight. A procedure slightly modified from the one described by (21) et al. was used for in-gel digestion. Briefly, the targeted protein bands from SDS-PAGE gel were manually excised, cut into 1 mm^3^pieces, and washed in 100 mM ammonium bicarbonate:acetonitrile (1:1, v/v) and 25 mM ammonium bicarbonate /acetonitrile (1:1, v/v), respectively to achieve complete destaining. The gel pieces were further treated with ACN, and dried *via* speed vac. After drying, gel pieces were reduced in 50 μl of 10 mM DTT at 56 °C for 60 min and then alkylated by 55 mM IAM for 30 min at room temperature. The gel pieces were washed with 25 mM ammonium bicarbonate:acetonitrile (1:1, v/v) twice to removed excess DTT and IAM. After drying, the gel pieces were placed on ice in 50 μL of trypsin solution at 10 ng/μL in 25 mM ammonium bicarbonate and incubated on ice for 60 min. Then, digestion was performed at 37 °C for 16 h. Peptide extraction was performed twice for 0.5 h with 100 μL 50% acetonitrile/0.2% formic acid. The combined extracts were concentrated in a Speed Vac to about 15 μL.

#### LC-MS/MS

Our mass spectrometry data were collected using an Orbitrap Fusion Lumos mass spectrometer (Thermo Fisher Scientific, San Jose, CA) coupled to a Eksigent Ekspert^TM^ nanoLC 425 System (Sciex). A Trap-Elute Jumper Chip (P/N:800-00389) and a coupled to a 1/16” 10 port Valco directed loading performed by the gradient 1 pump and final elution (by the gradient 2 pump). The column assembly was was designed as two tandem 75μmx15cm columns (ChromXP C18-CL, 3μm 120A, Eksigent part of AB SCIEX) mounted in the ekspert^TM^ cHiPLC system. For each injection, we loaded an estimated 0.5 ¼g of total digest. Peptides were separated in-line with the mass spectrometer using a 120 min gradient composed of linear and static segments wherein Buffer A is is 0.1% formic acid and B is 95%ACN, 0.1% Formic acid. The gradient begins first holds at 4% for 3 min then makes the following transitions (%B, min): (26, 48), (35, 58), (35, 64), (50, 72), (50, 78), (94, 84), (94, 96), (4, 100), (4, 120).

#### Tandem mass spectrometry on the LUMOS Orbitrap

Scan sequences began with a full survey (m/z 350 -1500) acquired on an Orbitrap Fusion Lumos mass spectrometer (Thermo) at a resolution of 60,000 in the off axis Orbitrap segment (MS1). Every 3 seconds of the gradient MS1 scans were acquired during the 120 min gradient described above. The most abundant precursors were selected among 2-8 charge state ions at a 2.0E5 threshold. Ions were dynamically excluded for 30 seconds if they were targeted twice in the prior 30 sec. Selected ions were isolated by a multi-segment quadrupole with a mass window on m/z 2, then sequentially subjected to both CID and HCD activation conditions in the IT and the ioin routing multipole respectively. The AGC target for CID was 4.0E04, 35% collision energy, an activation Q of 0.25 and a 100 milliseconds maximum fill time. Targeted precursors were also fragmented by high energy collision-induced dissociation (HCD) at 40% collision energy, and an activation Q of 0.25. HCD fragment ions were analyzed using the Orbitrap (AGC 1.2E05, maximum injection time 110 ms, and resolution set to 30,000 at 400 Th). Both MS2 channels were recorded as centroid and the MS1 survey scans were recorded in profile mode.

#### Proteomic Searches

Initial spectral searches were performed with Proteome Discoverer version 2.1.1.21 (Thermo) using Sequest HT. Spectra were also searched with Byonic search engine (Protein Metrics) ver. 2.8.2. Search databases were composed of the Uniprot KB for species 9606 (Human) downloaded 10/24/2016 containing 92645 sequences and Uniprot KB for taxonomy 562 (*E. coli*) downloaded on 11/08/2016 containing 10079 sequences. For Byonic searches, these two data bases were directly concatenated. In either search an equal number of decoy entries were created and searched simultaneously by reversing the original entries in the Target databases.

## ACKNOWLEDGEMENTS

This work was supported by a Cystic Fibrosis Foundation Pilot Award from the University of Iowa (R458-CR11). C.A.A. is supported by NIGMS (GM106569), is an American Heart Association Established Investigator (A22180002) and a member of the Membrane Protein Structural Dynamics Consortium (GM087519). We thank Dr. Julien Sebag for providing access to Spectramax i3.

